# Genome-scale identification of SARS-CoV-2 and pan-coronavirus host factor networks

**DOI:** 10.1101/2020.10.07.326462

**Authors:** William M. Schneider, Joseph M. Luna, H.-Heinrich Hoffmann, Francisco J. Sánchez-Rivera, Andrew A. Leal, Alison W. Ashbrook, Jérémie Le Pen, Eleftherios Michailidis, Inna Ricardo-Lax, Avery Peace, Ansgar F. Stenzel, Scott W. Lowe, Margaret R. MacDonald, Charles M. Rice, John T. Poirier

## Abstract

The COVID-19 pandemic has claimed the lives of more than one million people worldwide. The causative agent, SARS-CoV-2, is a member of the *Coronaviridae* family, which are viruses that cause respiratory infections of varying severity. The cellular host factors and pathways co-opted by SARS-CoV-2 and other coronaviruses in the execution of their life cycles remain ill-defined. To develop an extensive compendium of host factors required for infection by SARS-CoV-2 and three seasonal coronaviruses (HCoV-OC43, HCoV-NL63, and HCoV-229E), we performed parallel genome-scale CRISPR knockout screens. These screens uncovered multiple host factors and pathways with pan-coronavirus and virus-specific functional roles, including major dependency on glycosaminoglycan biosynthesis, SREBP signaling, and glycosylphosphatidylinositol biosynthesis, as well as an unexpected requirement for several poorly characterized proteins. We identified an absolute requirement for the VTT-domain containing protein TMEM41B for infection by SARS-CoV-2 and all other coronaviruses. This human *Coronaviridae* host factor compendium represents a rich resource to develop new therapeutic strategies for acute COVID-19 and potential future coronavirus spillover events.

**HIGHLIGHTS:** Genome-wide CRISPR screens for SARS-CoV-2, HCoV-OC43, HCoV-NL63, and HCoV-229E coronavirus host factors.

Parallel genome-wide CRISPR screening uncovered host factors and pathways with pan-coronavirus and virus-specific functional roles.

Coronaviruses co-opt multiple biological pathways, including glycosaminoglycan biosynthesis, SREBP signaling, and glycosylphosphatidylinositol biosynthesis and anchoring, among others.

TMEM41B - a poorly understood factor with roles in autophagy and lipid mobilization - is a critical pan-coronavirus host factor.

## INTRODUCTION

Severe acute respiratory syndrome coronavirus 2 (SARS-CoV-2), the causative agent of the ongoing coronavirus disease 2019 (COVID-19) pandemic, has claimed the lives of more than one million people worldwide in a span of nine months (Zhou et al., 2020; Zhu et al., 2020a), (https://coronavirus.jhu.edu/map.html). SARS-CoV-2 is a betacoronavirus from the *Coronaviridae* family, which is composed of enveloped positive-sense RNA viruses with large (> 30kb) genomes that can infect a variety of vertebrate hosts (Cui et al., 2019). Seasonal human coronaviruses (HCoVs), such as the HCoV-OC43 betacoronavirus, as well as the HCoV-NL63 and HCoV-229E alphacoronaviruses, can cause mild to moderate upper-respiratory infections with cold-like symptoms in humans (Cui et al., 2019). In stark contrast, highly pathogenic betacoronaviruses have been responsible for multiple deadly outbreaks in the 21st century, including severe acute respiratory syndrome coronavirus (SARS-CoV, 2003), Middle East respiratory syndrome coronavirus (MERS-CoV, 2012), and SARS-CoV-2 (2019) (Cui et al., 2019). The spread of SARS-CoV and MERS-CoV was contained, in part due to their comparatively low transmissibility (Cui et al., 2019). However, SARS-CoV-2 spreads more readily and remains largely uncontrolled across the globe, presenting an urgent health crisis.

A complete understanding of the host factors and pathways co-opted by SARS-CoV-2 and other coronaviruses for the execution of their life cycles could contribute to the development of therapies to treat COVID-19 and increase preparedness for potential future outbreaks. Large scale forward genetic approaches based on RNA interference, insertional mutagenesis, and CRISPR have proven powerful for identifying host factors required for infection by different viruses (reviewed in (Puschnik et al., 2017)). Here, we performed parallel genome-scale CRISPR-Cas9 knockout screens to generate an extensive functional catalog of host factors required for infection by SARS-CoV-2 and three seasonal coronaviruses (HCoV-OC43, HCoV-NL63, and HCoV-229E). We identified multiple genes and pathways with pan-coronavirus and virus-specific functional roles, including factors involved in glycosaminoglycan (GAG) biosynthesis, sterol regulatory element-binding protein (SREBP) signaling, and glycosylphosphatidylinositol (GPI) biosynthesis, as well as several poorly characterized proteins, such as Transmembrane Protein 41B (TMEM41B). We show that the <u>V</u>MP1, <u>T</u>MEM41, and <u>T</u>VP38 (VTT)-domain containing protein TMEM41B is a critical host factor required for infection by SARS-CoV-2, HCoV-OC43, and HCoV-229E, as well as several flaviviruses of high interest to public health (Hoffmann & Schneider et al.,; see accompanying manuscript), thereby nominating it as a broad-spectrum RNA virus liability and potential high-priority target for future drug development efforts.

## RESULTS

### Genome-wide CRISPR screens identify host factors required for SARS-CoV-2 infection

We set out to develop an extensive catalog of human host factors required for infection by the pandemic SARS-CoV-2 strain and three seasonal coronaviruses (HCoV-OC43, HCoV-NL63 and HCoV-229E) (Cui et al., 2019) (**Figure 1A**). We employed high-throughput CRISPR-Cas9 genetic screening using the well-established Brunello genome-wide library, which is composed of 76,441 single guide RNAs (sgRNAs) targeting 19,114 human genes (Doench et al., 2016a). As a screening platform, we used Cas9-expressing Huh-7.5 hepatoma cells (Huh-7.5-Cas9), which endogenously express the SARS-CoV-2 cellular receptor, angiotensin-converting enzyme 2 (ACE2), as well as transmembrane serine protease 2 (TMPRSS2), a key mediator of SARS-CoV-2 entry (Hoffmann et al., 2020b). We recently showed that Huh-7.5-Cas9 cells are permissive to infection by SARS-CoV-2, HCoV-OC43, HCoV-NL63, and HCoV-229E, and that they are a robust system for CRISPR-based genetic screening (Hoffmann et al., 2020a).

**Figure 1.**
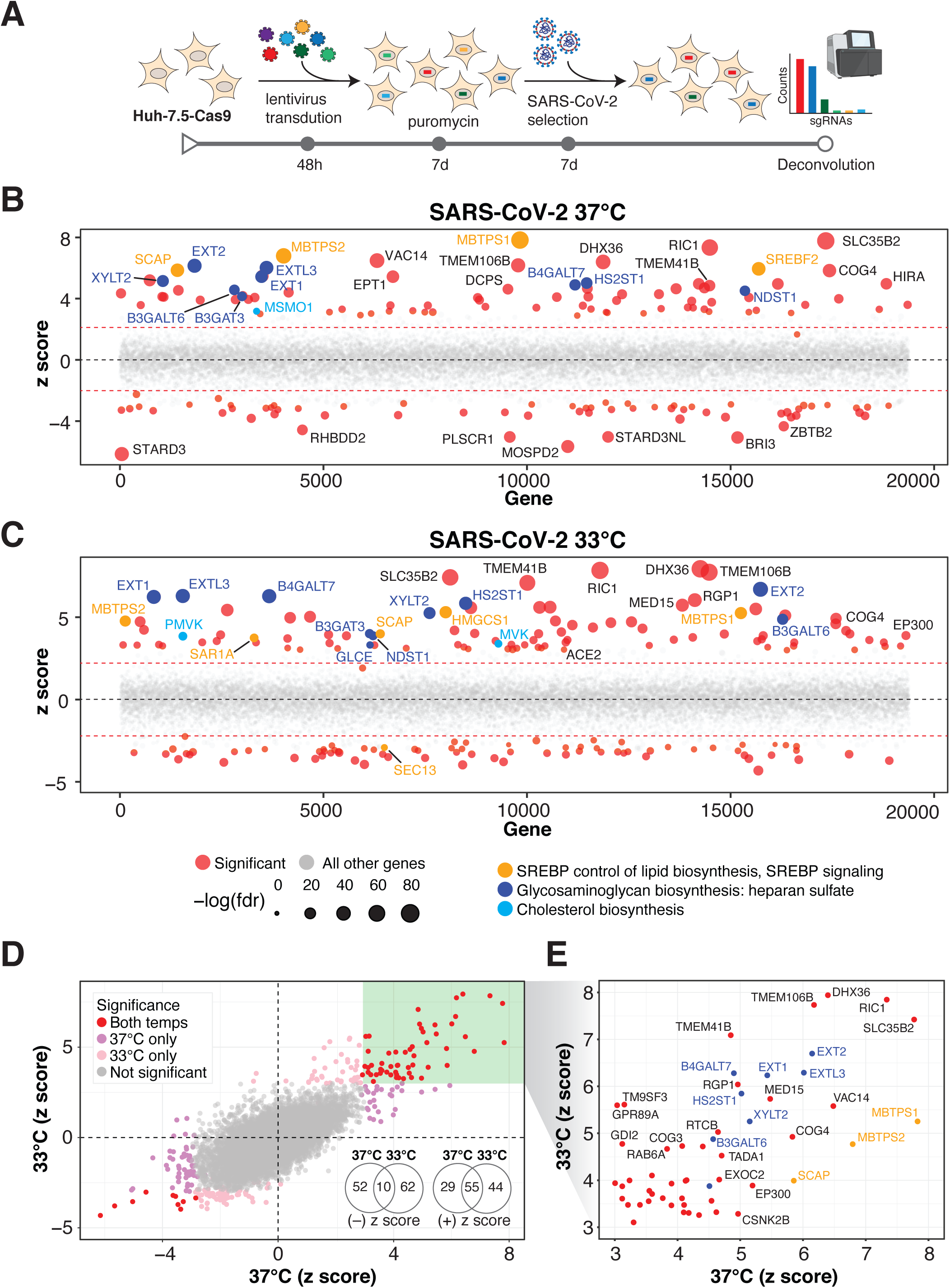
Genome-wide CRISPR screens identify host factors required for SARS-CoV-2 infection. (A) Genome-wide CRISPR screening workflow. Cas9-expressing Huh-7.5 cells are transduced with the Brunello genome-wide CRISPR library, selected with puromycin, and infected with SARS-CoV-2 or one of three season coronaviruses (HCoV-OC43, HCoV-NL63, or HCoV-229E). Surviving cells along with mock controls are then harvested and sgRNA abundance is determined using next-generation sequencing. (B) Bubble plot of data from SARS-CoV-2 screens at 37 °C. Red lines denote z = ± 2. (C) Bubble plot of data from SARS-CoV-2 screens at 33 °C. Red lines denote z = ± 2. (D) Scatterplot comparing z-scores from (B) and (C) for SARS-CoV-2 screens at 37 °C and 33 °C, respectively. (E) Subset of significantly enriched genes from SARS-CoV-2 screens at 37 °C and 33 °C.

We performed a series of parallel genetic screens by transducing Huh-7.5-Cas9 cells with the Brunello sgRNA library followed by antibiotic selection and expansion for seven days to ensure homogeneous CRISPR-based knockout of host factor genes prior to infection with the panel of coronaviruses. In this context, cells expressing sgRNAs targeting genes required for virus infection or virus-induced death should survive while those expressing neutral sgRNAs or sgRNAs targeting genes irrelevant to infection are expected to deplete. Similarly, cells expressing sgRNAs targeting essential genes with no roles in virus infection or virus-induced death are expected to deplete in both “mock” (uninfected) and virus-infected conditions. SARS-CoV-2 screens were performed in triplicate at two physiologically relevant temperatures, 33 °C and 37 °C, to mimic the temperatures of the upper and lower airways, respectively (V’kovski et al., 2020). Surviving cells were harvested five days post-infection and subjected to genomic DNA extraction and screen deconvolution using high-throughput sequencing (Hoffmann et al., 2020a).

Several quality control (QC) metrics demonstrated excellent technical performance across all screens and biological replicates (**Figure S1**). First, we confirmed that 76,160/76,441 (99.6%) of sgRNAs from the Brunello library were recovered from the plasmid preparation and that all screen libraries were sequenced to saturation (**Figure S1A**). Second, pairwise correlation analyses demonstrated that biological replicates from each genetic screen clustered together and shared a high correlation coefficient (**Figure S1B)**. Third, receiver operating characteristic (ROC) curves generated based on the fitness effects of disruption of previously defined neutral and essential genes from the Brunello library confirmed robust gene disruption in the cell pools (**Figure S1C-D**). The HCoV-229E screen, though successful, was particularly stringent, resulting in a lower area under the curve (AUC) relative to the other screens in this study. As described in our recent study (Hoffmann et al., 2020a), we performed a z-score analysis as well as gene essentiality (beta) score using a published maximum likelihood estimation (MLE) algorithm (Li et al., 2014). The gene essentiality analysis allowed us to stratify candidate host factor targets based on their effects on cellular fitness under mock (uninfected) conditions followed by the identification of high-confidence gene hits in virus-infected cells. Specifically, genes with beta scores similar to essential genes could affect cell survival in the presence or absence of infection and may be confounded by effects on cellular fitness. Conversely, genes with beta scores similar to neutral sgRNAs are predicted to affect cell survival only during viral infection and are more likely to be true positives. We implemented these parameters into our analysis pipeline and identified 146 and 171 genes that significantly influenced SARS-CoV-2-induced cell death at 37 °C and 33 °C, respectively (FDR < 0.05) (summarized in **Figure 1B-E, Figure S2B-C**, and **Table S1A-B**). These included the SARS-CoV-2 receptor ACE2 and the well known host factor cathepsin L (CTSL) (Hoffmann et al., 2020b; Letko et al., 2020; Yeager et al., 1992). MERS-CoV receptor DPP4 (Earnest et al., 2017; Wang et al., 2013) and putative SARS-CoV-2 receptors KREMEN1 and ASGR1 did not score in any of these screens (Gu et al., 2020). (**Figure S2A**). A total of 84 (37 °C) and 99 (33 °C) genes scored as candidate host factors that may facilitate SARS-CoV-2 infection (z-score > 0; FDR < 0.05). Conversely, 62 (37 °C) and 72 (33 °C) genes scored as candidate antiviral host factors (z-score < 0; FDR < 0.05). As expected, neutral and essential gene-targeting sgRNAs scored similarly across mock and SARS-CoV-2 conditions (blue and red dots in **Figure S2B-C**, respectively). Integrating the 33 °C and 37 °C SARS-CoV-2 screening datasets allowed us to obtain a clearer picture of candidate temperature-specific host factors that either support or antagonize SARS-CoV-2 viral infection (**Figure 1D-E** and **Figure S2D**). These results demonstrate that the human genome encodes a catalog of host factors that functionally contribute to the SARS-CoV-2 life cycle.

### Parallel genome-wide CRISPR screening against multiple human coronaviruses uncovers host factor networks with pan-coronavirus and virus-specific functional roles

Viruses within the same family often require the same host factors to complete their respective life cycles (Dimitrov, 2004). Nevertheless, there are several examples of closely related viruses with discrete host factor requirements. For example, while SARS-CoV-2 and HCoV-NL63 both engage ACE2 as a cellular receptor, HCoV-229E uses ANPEP while HCoV-OC43 has no known essential proteinaceous receptor (Cui et al., 2019; Forni et al., 2017). Beyond attachment factors and receptors, closely related viruses can also exploit specific components of multiple intracellular pathways in a virus-specific manner. For example, we recently reported that SARS-CoV-2 appears to rely on the RAB10 and RAB14 small GTPases whereas HCoV-229E, HCoV-OC43, and HCoV-NL63 mainly require RAB2A and RAB7A (Hoffmann et al., 2020a). A comprehensive functional understanding of the commonalities and differences among coronaviruses and other virus families could pave the way for both specific and general antiviral therapies. Towards this goal, we significantly expanded our functional genomics efforts to develop an extensive functional catalog of human host factors required for infection by members of the *Coronaviridae* family, including two alphacoronaviruses (HCoV-NL63 and HCoV-229E) and one additional betacoronavirus (HCoV-OC43).

The results of these three additional CRISPR screens are shown in **Figure 2A-C, Figure S3** and **Table S1C-E**. An integrative analysis that also includes the two SARS-CoV-2 screens described above is shown in **Figure 2D**. Taken together, these screens identified numerous coronavirus-specific and pan-coronavirus host factors that appear to play critical roles during infection by each of these viruses. This extensive network of human host factors functionally implicates numerous cellular pathways, as shown in **Figure 3A**. We present a selection of comparative analyses below that highlight both pan-coronavirus and virus-specific host factors through the lens of SARS-CoV-2.

**Figure 2.**
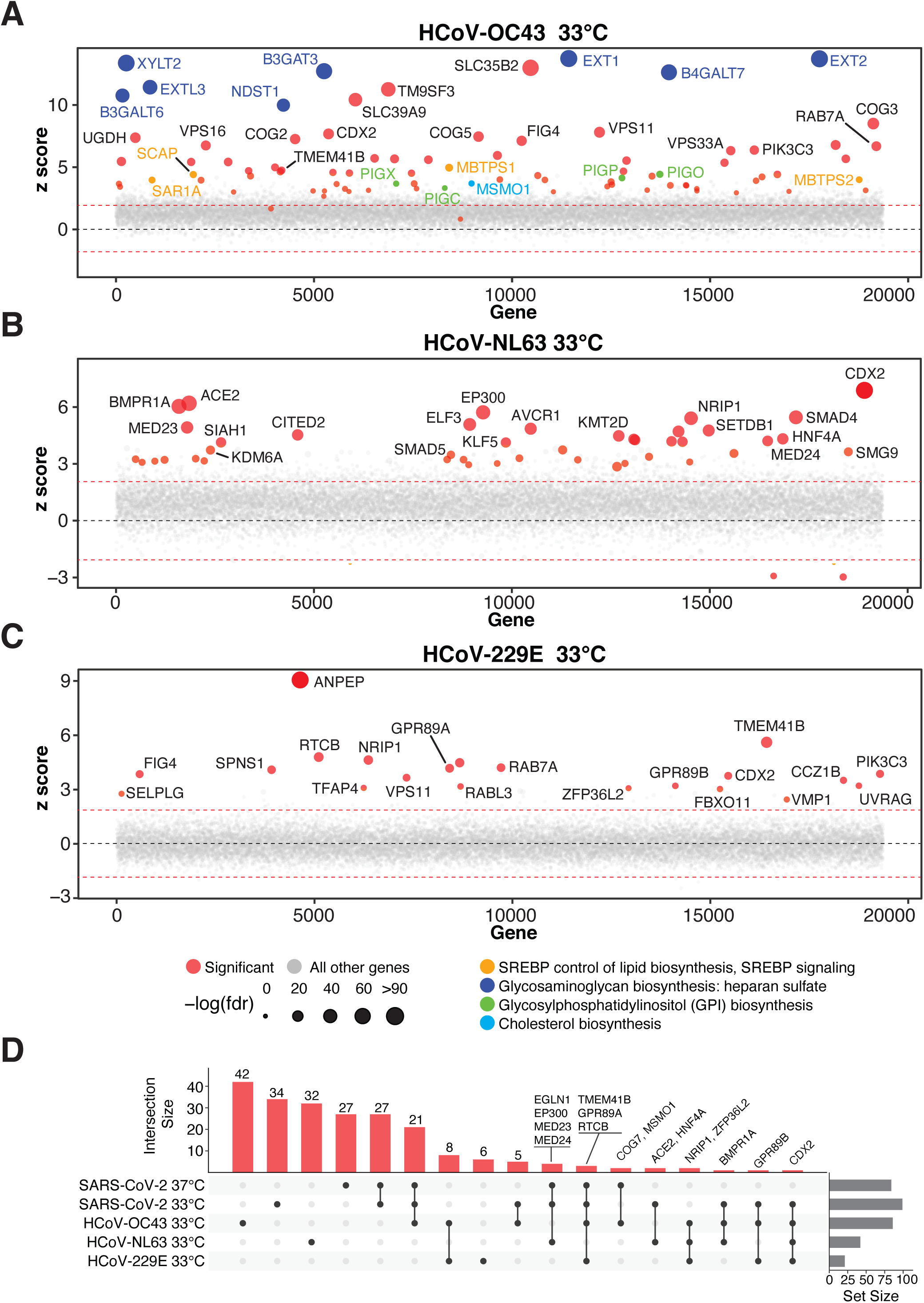
Parallel genome-wide CRISPR screening against multiple human coronaviruses uncovers host factors and pathways with pan-coronavirus and virus-specific functional roles. (A) Bubble plot of data from HCoV-OC43 screens at 33 °C. Red lines denote z = ±2. (B) Bubble plot of data from HCoV-NL63 screens at 33 °C. Red lines denote z = ±2. (C) Bubble plot of data from HCoV-229E screens at 33 °C. Red lines denote z = ±2. (D) UpSet plot showing enriched hits overlapping in screens across all four viruses. Select genes for enriched sgRNAs are indicated.

**Figure 3.**
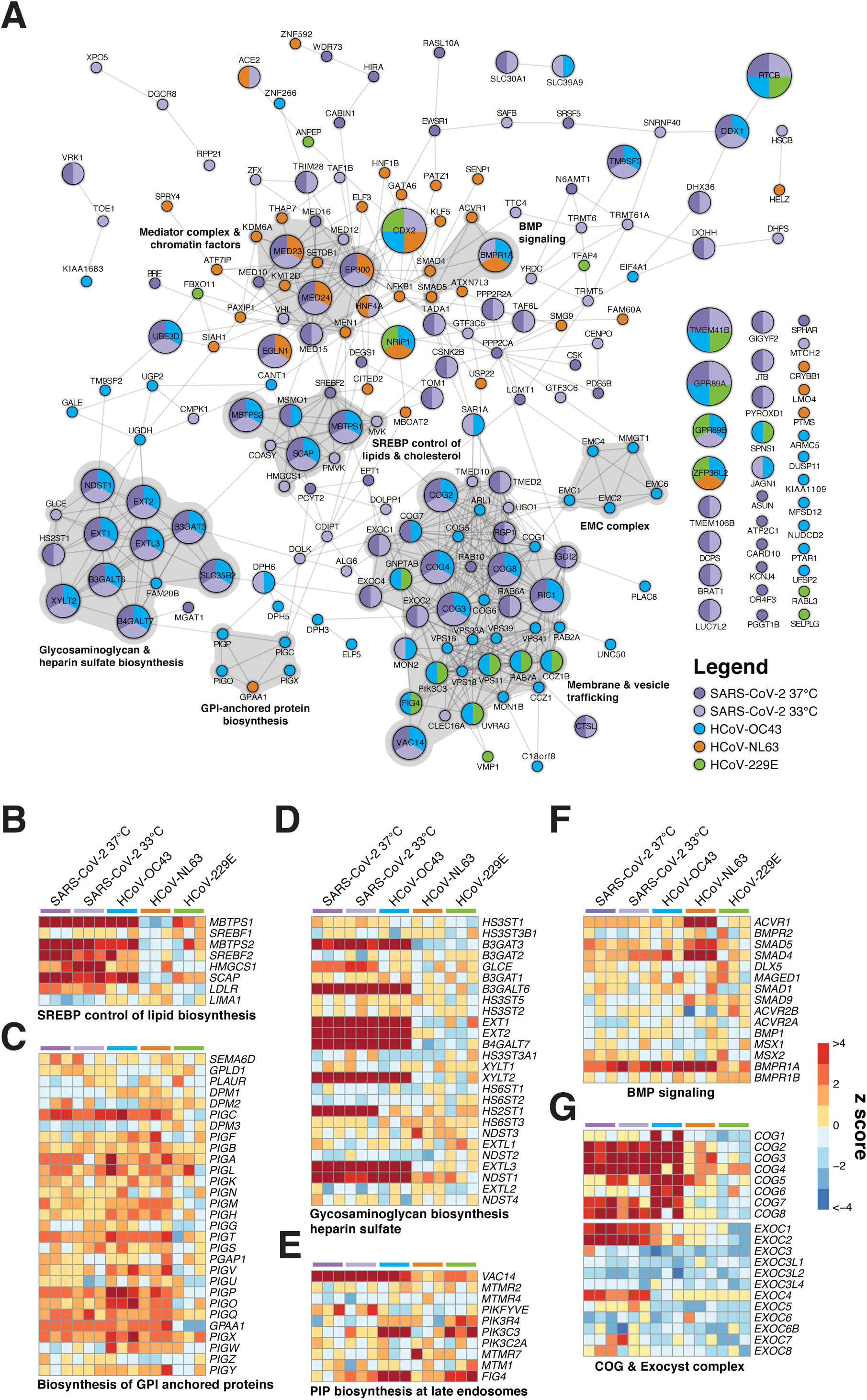
Coronaviruses co-opt an extensive network of human proteins and pathways to complete their life cycle. (A) Network analysis of human coronavirus host factors for all significant screening hits using the STRING-db protein:protein interaction network. Nodes are subdivided by number of virus screens by color and size for which the node was significant. Highly interconnected and functionally related genes are further highlighted in gray. (B-G) Comparative pathway-focused heatmaps showing enriched and depleted sgRNAs across all CRISPR screens.

### SARS-CoV-2 and HCoV-OC43 (betacoronaviruses)

The SARS-CoV-2 and HCoV-OC43 betacoronaviruses co-opt a significantly overlapping set of host factors to carry out their life cycles. These include proteins involved in pathways related to GAG biosynthesis (e.g., heparan sulfate) and transport, such as EXT1, EXT2, EXTL3, B3GALT6, B3GAT3, B4GALT7, SLC35B2, XYLT2, HS2ST1, and NDST1 (Aikawa et al., 2001; Bai et al., 2001; Casanova et al., 2008; Cuellar et al., 2007; Kitagawa et al., 1998; Kreuger and Kjellén, 2012; Lind et al., 1998; Okajima et al., 1999; Ponighaus et al., 2007). We also identified multiple factors that regulate intracellular protein trafficking, processing, and sorting through the cis-oligomeric Golgi (COG) complex, including COG2, COG3, COG4, COG7, and COG8 (Blackburn et al., 2019; Smith and Lupashin, 2008) (**Figure 3A**,**D**,**G**). Consistent with the role of heparan sulfate as an attachment factor for multiple viruses, the heparan sulfate biosynthesis pathway has been previously implicated as a critical host pathway for several viruses and virus families, including herpes simplex virus (O’Donnell and Shukla, 2008), human papillomavirus (Giroglou et al., 2001; Joyce et al., 1999), respiratory syncytial virus (Bourgeois et al., 1998; Escribano-Romero et al., 2004; Feldman et al., 2000; Hallak et al., 2000; Harris and Werling, 2003; Karger et al., 2001; Krusat and Streckert, 1997; Martinez and Melero, 2000; Techaarpornkul et al., 2002), adenoviruses (Dechecchi et al., 2001; Dechecchi et al., 2000), hepatitis C virus (Xu et al., 2015), dengue and Zika virus (Cruz-Oliveira et al., 2015; Marceau et al., 2016; Savidis et al., 2016), West Nile virus (Perera-Lecoin et al., 2013), Rift Valley fever virus (Riblett et al., 2016), Eastern equine encephalitis virus (Gardner et al., 2011), and HIV (Ibrahim et al., 1999), among others. These studies support the central functional role of heparan sulfate proteoglycans and other glycosaminoglycans as common mediators of binding and entry for many viruses. Indeed, recent cellular and biochemical evidence suggests that SARS-CoV-2 exploits heparan sulfate proteoglycans cooperatively with ACE2 to bind to and gain entry into cells (Clausen et al., 2020b). Given that no protein has been identified as a cellular receptor for HCoV-OC43 entry, it is possible that this betacoronavirus engages one or more GAGs to invade target cells.

Another set of top-scoring factors in common between the SARS-CoV-2 and HCoV-OC43 screens were related to cholesterol homeostasis, particularly those related to SREBP cleavage-activating protein (SCAP)-mediated cholesterol sensing and the SREBP pathway (**Figure 3A-B, Figure S4A-B**). Indeed, genes known to be functionally involved in the sensing and biosynthesis of cholesterol, such as SCAP, SREBF2 (but not SREBF1), MBTPS1, MBTPS2, and SAR1A were among the top enriched genes for these two viruses (**Figure 3A-B**). These results are consistent with our recent discovery that SARS-CoV-2 and other coronaviruses require SCAP, NPC2, and EMC1 to carry out their life cycle (Hoffmann et al., 2020a) and extend these findings to further elaborate the essential regulatory components of SREBP signaling. We also note a unique reliance by HCoV-OC43 on factors involved in the synthesis of GPI anchored proteins (**Figure 3A**,**C)**. Collectively, these results nominate factors involved in GAG biosynthesis and transport, intracellular protein trafficking, processing, and sorting, and cholesterol homeostasis as potential targetable factors to inhibit SARS-CoV-2 and HCoV-OC43.

### SARS-CoV-2-specific host factors

We also identified host factors and pathways that appear to be required for SARS-CoV-2 infection but less so for other coronaviruses tested. Genes that participate in the mevalonate pathway, which is regulated by SREBP and is responsible for converting mevalonate into sterol isoprenoids, such as cholesterol (Buhaescu and Izzedine, 2007; Goldstein and Brown, 1990), were among the top scoring hits (**Figure 3A**,**B, Figure S4A, Figure S5A**). Multiple sgRNAs targeting 3-Hydroxy-3-Methylglutaryl-CoA Synthase 1 (*HMGCS1*), which catalyzes the conversion of HMG-CoA to mevalonic acid, mevalonate kinase (*MVK*), which catalyzes the phosphorylation of mevalonic acid into phosphomevalonate, and phosphomevalonate kinase (*PMVK*), which converts phosphomevalonate to mevalonate 5-diphosphate, were significantly enriched in SARS-CoV-2 screens, although to a lesser extent than SREBP signaling (**Figure 3A**,**B** and **Figure S5A**). These results suggest that factors and intermediates of the mevalonate pathway, which are known to play important roles in post-translational modification of many proteins involved in key processes, such as intracellular signaling and protein glycosylation (Buhaescu and Izzedine, 2007; Goldstein and Brown, 1990), are important for the life cycle of SARS-CoV-2.

Another set of top-scoring genes in SARS-CoV-2 screens encode multiple subunits of the exocyst complex, which regulates the tethering of secretory vesicles to the plasma membrane and their subsequent SNARE-mediated membrane fusion and exocytosis (Martin-Urdiroz et al., 2016; Wu and Guo, 2015) (**Figure 3G**). Multiple sgRNAs targeting *EXOC1, EXOC2*, and *EXOC4* were significantly enriched, suggesting a critical role for these factors in mediating SARS-CoV-2 infection and virus-induced cell death. The mammalian exocyst complex is known to interact with specific RAB GTPases to coordinate intracellular trafficking (Babbey et al., 2010; Mei and Guo, 2018). Indeed, sgRNAs targeting RAB6A and RAB10 were also among the most significantly enriched hits in SARS-CoV-2 screens (**Figure 3**), a finding that is consistent with our recent work identifying RAB10 as a putative SARS-CoV-2-specific host factor (Hoffmann et al., 2020a). These results suggest that SARS-CoV-2 relies on specific intracellular host factors and complexes that govern intracellular transport.

Another complex that appears to play an essential role in the SARS-CoV-2 life cycle is the Mediator complex (**Figure 3** and **Figure S5B**). The mammalian Mediator is an evolutionarily conserved protein complex composed of at least 28 subunits that regulates transcription by functionally connecting general transcription factors with the core transcriptional machinery (Allen and Taatjes, 2015). The Mediator subunits MED10, MED12, MED15, MED16, MED23, and MED24 were among the top-scoring genes in SARS-CoV-2 screens, suggesting a critical role for this complex during infection and death by this virus (**Figure 3**). Intriguingly, a non-overlapping set of Mediator subunits was recently implicated in HIV-1 replication, including MED6, MED7, MED11, MED14, MED21, MED26, MED27, MED28, and MED30 (Ruiz et al., 2014), suggesting that different viruses might have specific requirements for members of this complex during transcription and replication of their genomes.

Beyond well-characterized pathways that were represented by multiple components, we also identified factors with less understood network-level connections (**Figure 3A**). These include the EMC genes (**Figure S5C)**, DEAH-Box helicases *DHX36* and *DHX38*, Golgin family proteins *GOLGA6L1* and *GOLGA8O*, the General Transcription Factor IIIC subunits *GTF3C5* and *GTF3C6*, tRNA methyltransferases *TRMT5* and *TRMT6*, G protein-coupled receptors *GPR89A* and *GPR89B*, Transmembrane P24 trafficking proteins *TMED2* and *TMED10*, and genes involved in phosphatidylethanolamine biosynthesis, such as *PCYT2* and *EPT1*, among others (**Figure 3A** and **Table S1**). Further mining of SARS-CoV-2 host factor networks could expand the repertoire of potential targetable factors to treat COVID-19.

Given the genome-scale depth of this screening data, we sought to determine if interactome focused networks were significantly enriched in a genome-wide context. A recent SARS-CoV-2 CRISPR screen in an African green monkey Vero-E6 cells (Wei et al., 2020) failed to detect significant enrichment from a SARS-CoV-2-human protein-protein network derived from IP-MS (Gordon et al., 2020). We recently tested genes from this protein-protein interactome with a focused CRISPR screen and assigned functional relevance to numerous members (Hoffmann et al., 2020b). In our genome scale screens, we subsetted z-scores from the full interactome in Gordon et al., and detected a modest significant enrichment for hits in a genome-wide context (**Figure S5D**). Upon subsetting the functionally relevant members from our focused screen in Hoffmann et al., we observed a striking increase in significance (**Figure S5D**). The subsetted heatmap of hits in Hoffmann et al., mapped onto genome scale data largely cross validated many, but not all of our prior key findings (**Figure S5E**). These results demonstrate the power and utility of focused CRISPR screens to complement the breadth of genome-wide efforts.

### HCoV-NL63 and HCoV-229E (alphacoronaviruses)

Interestingly, the catalog of host factors essential for infection by the HCoV-NL63 and HCoV-229E alphacoronaviruses is significantly different than that of betacoronaviruses (**Figure 2, Figure 3A**, and **Figure S4C-D**). Furthermore, these viruses apparently rely on substantially different pathways to carry out their life cycles. Consistent with the literature, both screens successfully identified the cognate virus receptors, as evidenced by robust enrichment of sgRNAs targeting ACE2 (HCoV-NL63) and ANPEP (HCoV-229E) (Hoffmann et al., 2020b; Letko et al., 2020; Yeager et al., 1992) (**Figure 2** and **Figure 3**). Intriguingly, HCoV-NL63 seems to rely on a core set of host chromatin regulators with known functional interactions, including EP300, KDM6A (also known as UTX), KMT2D (also known as MLL4), MED23, MED24, MEN1, PAXIP1, and SETDB1. This raises the tantalizing possibility that HCoV-NL63 co-opts the well established UTX-MLL4-EP300 enhancer remodeling network (which also contains PAXIP1) (Wang et al., 2017) to reprogram the host transcriptome for successful infection (**Figure 3A orange nodes**). In addition, we observed a requirement for factors involved in BMP signaling with HCoV-NL63 specific factors SMAD4, SMAD5, ACVR1, and BMPR1A (**Figure 3F)**. Lastly, our screening results suggest that HCoV-229E is particularly dependent on members of the HOPS complex, including VPS11 and RAB7A (**Figure 3A**) (Balderhaar and Ungermann, 2013). Collectively, our results demonstrate that even closely related viruses, such as the HCoV-NL63 and HCoV-229E alphacoronaviruses, can employ markedly different host factor pathways during infection.

### Integrative analyses identify TMEM41B as a critical host factor for SARS-CoV-2

As shown in **Figure 1B-E, Figure 2**, and **Figure 3A**, sgRNAs targeting TMEM41B were among the most significantly enriched sgRNAs across the SARS-CoV-2, HCoV-OC43, and HCoV-229E screens. TMEM41B is a poorly understood ER-localized transmembrane protein that was recently implicated in the autophagy pathway (Moretti et al., 2018; Morita et al., 2018; Shoemaker et al., 2019). Specifically, TMEM41B deficiency was shown to lead to accumulation of ATG proteins, thereby blocking the autophagy pathway at the early step of autophagosome formation. In addition, TMEM41B deficiency was shown to trigger the abnormal accumulation of intracellular lipid droplets. These phenotypes have been linked to the function of another autophagy factor, vacuole membrane protein 1 (VMP1), which shares a rare and characteristic VTT domain (Morita et al., 2019). Interestingly, multiple sgRNAs targeting VMP1 were significantly enriched in the HCoV-229E screen (**Figure 2C**) while there was a trend towards enrichment in the SARS-CoV-2 screens that did not meet the cutoff for statistical significance due to variability between replicates. These findings prompted us to look into the autophagy network across all coronavirus screens in more detail.

TMEM41B was the only autophagy-related gene that scored as a significant hit across our multiple coronavirus screens while only a handful of genes involved in the nucleation and tethering steps scored for HCoV-OC43 and HCoV-299E (**Figure 4A**). This was a striking finding given that we concurrently identified TMEM41B as a critical host factor for infection by multiple biosafety level three (BSL3) and BSL4 members of the *Flaviviridae* family, including 1) the mosquito-borne flaviviruses Zika virus, yellow fever virus, West Nile virus, and dengue virus, 2) the tick-borne flaviviruses Powassan virus, tick-borne encephalitis virus, Kyasanur forest disease virus, Omsk hemorrhagic fever virus, and Alkhurma hemorrhagic fever virus, and 3) other members of the *Flaviviridae*, including hepatitis C virus from the hepacivirus genus and bovine viral diarrhea virus from the pestivirus genus (Hoffmann & Schneider et al.,; see accompanying manuscript). Comparative analyses between one such flavivirus, Zika, and SARS-CoV-2 screens suggested that TMEM41B is the only host factor that is critical for infection by both viruses (**Figure 4B**) (Hoffmann & Schneider et al.,; see accompanying manuscript). Indeed, genetic disruption of TMEM41B was sufficient to render cells resistant to infection by SARS-CoV-2 and all of the seasonal coronaviruses and, importantly, infectivity could be reconstituted by TMEM41B cDNA expression (**Figure 4C**). Interestingly, TMEM41B sub-cellular localization changed dramatically upon infection by SARS-CoV-2 (**Figure 4D**). While mock-infected cells exhibited a diffuse reticular-like pattern of TMEM41B localization in the cytoplasm, cells infected with SARS-CoV-2 were characterized by large TMEM41B cytosolic aggregates and positive immunostaining for SARS-CoV-2 NS4A and NS4B viral proteins (**Figure 4D**). Thus, TMEM41B is a critical host factor required for infection by coronaviruses and several other pathogenic viruses and represents an attractive target for further intensive study.

**Figure 4.**
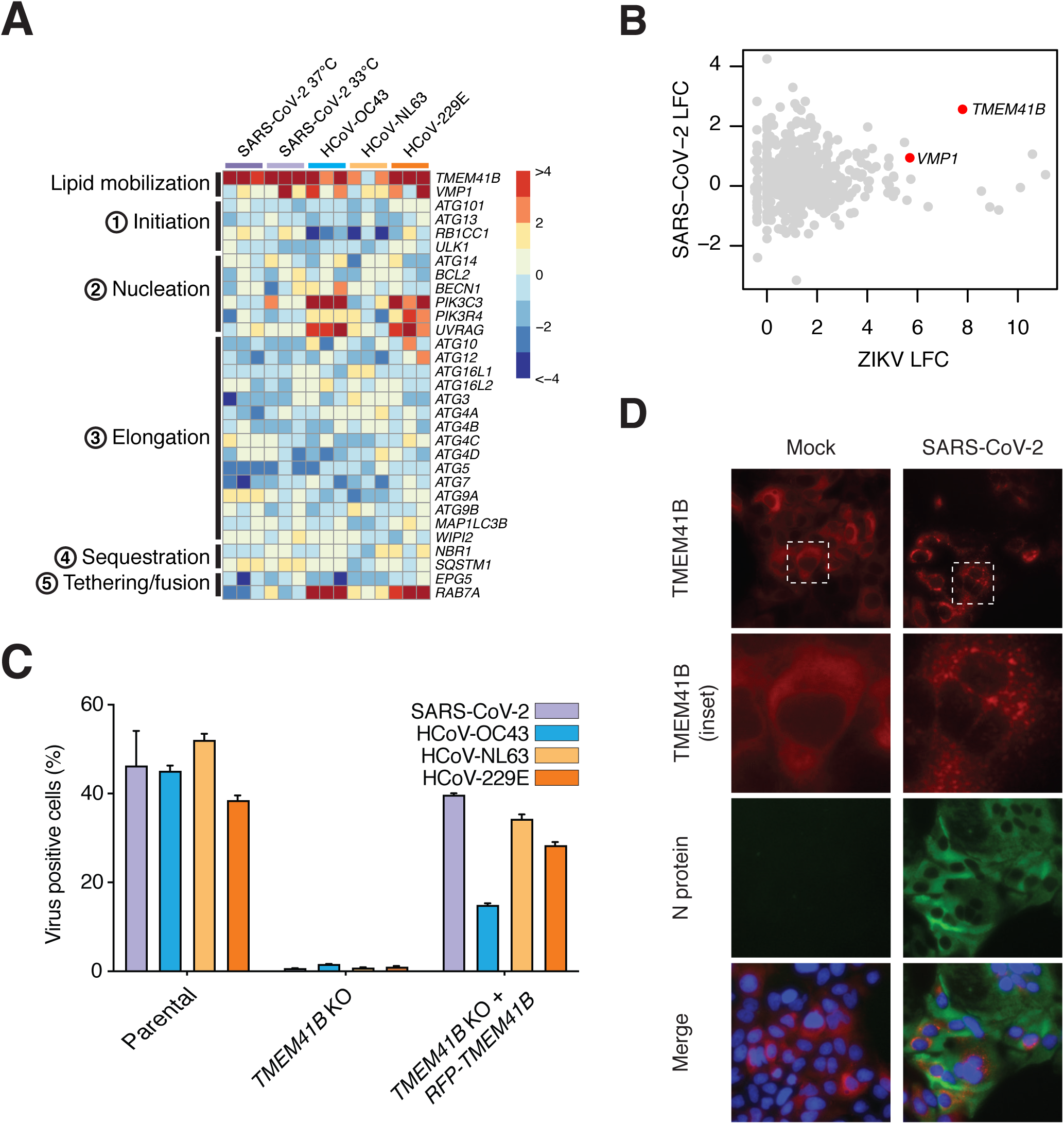
Integrative analyses identify TMEM41B as a critical host factor for SARS-CoV-2, HCoV-OC43, and HCoV-229E. (A) Heatmap of genes from the autophagy pathway ordered sequentially by function in the autophagy cascade. (B) Comparative analysis of Zika and SARS-CoV-2 screens by log fold change (LFC). (C) Coronavirus infectivity assay in Huh-7.5 parental cells, TMEM41B knockout cells, and TMEM41B knockout cells reconstituted with TMEM41B cDNA. Infections were carried out under the following optimized conditions: SARS-CoV-2 (500 PFU; 48 h) = MOI of 0.05 PFU/cell; HCoV-OC43 (15,000 PFU; 72 h) = MOI = 2 PFU/cell; HCoV-NL63 (500 PFU; 72 h) = MOI = 0.05 PFU/cell; HCoV-229E (2,500 PFU; 24 h) = 0.15 PFU/cell. (D) Fluorescence microscopy images of SARS-CoV-2 immunostaining (green), TMEM41B-mCherry (red), and DNA (DAPI).

## DISCUSSION

The full complement of human proteins and pathways required for infection by SARS-CoV-2 remains poorly defined. A more complete understanding of the cellular and molecular mechanisms that are co-opted by coronaviruses could catalyze drug development efforts to combat the ongoing COVID-19 pandemic and prepare for potential future outbreaks. We performed parallel genome-scale CRISPR-Cas9 knockout screens to generate the most extensive functional catalog of host factors required for infection by SARS-CoV-2 and three seasonal coronaviruses (HCoV-OC43, HCoV-NL63, and HCoV-229E) reported to date (**Figure 3A** and **Table S1**). This catalog contains multiple host factors and pathways that play critical pan-coronavirus and virus-specific functional roles (**Figure 3A**).

We identified a profound requirement for factors involved in GAG (e.g., heparan sulfate) biosynthesis, transport, and modification during infection by the SARS-CoV-2 and HCoV-OC43 betacoronaviruses. These results suggest that HCoV-OC43 might engage one or more GAGs to invade target cells, and support recent studies showing that SARS-CoV-2 simultaneously co-opts heparan sulfate proteoglycans and ACE2 to gain entry into cells (Clausen et al., 2020a). Of note, HS2ST1, a component of the heparan sulfate biosynthetic pathway, was identified in the present study as well as in our prior work (Hoffmann et al., 2020a). These results nominate factors involved in GAG biosynthesis, modification, and transport as potential targetable factors to inhibit SARS-CoV-2 and HCoV-OC43.

We also identified proteins that mediate cholesterol homeostasis via SREBP signaling as key host factors for infection by SARS-CoV-2 and HCoV-OC43. These results agree with and substantially extend results from our recent functional interactome study demonstrating that SARS-CoV-2 and other coronaviruses require SCAP, NPC2, and EMC1 to carry out their life cycles and elucidate the essential regulatory components of SREBP signaling (Hoffmann et al., 2020a). In addition, these results are consistent with work on several other viruses, including Ebola virus (Carette et al., 2011) and hantavirus (Kleinfelter et al., 2015), among others. Beyond SREBP signaling, we also identified three of the key enzymes from the mevalonate pathway (HMGCS1, MVK, and PMVK), which is responsible for converting mevalonate into sterol isoprenoids, such as cholesterol (**Figure 3A, Figure 3B, Figure S4A**). Given that statins – one of the most widely used class of drugs in the world – block mevalonate production via inhibition of the HMGCoA reductase and have been recently associated with improved outcomes among COVID-19 patients (Fajgenbaum and Rader, 2020; Zhang et al., 2020), it is tempting to speculate that pharmacological modulation of the mevalonate pathway could be a promising strategy for treating COVID-19.

Another set of critical host factors for SARS-CoV-2 infection are involved in regulating intracellular transport, tethering, and exocytosis of secretory vesicles, including RAB GTPases (**Figure 3A, Figure S4E**). These results lend further support to our recent nomination of RAB10 as a putative SARS-CoV-2 host factor (Hoffmann et al., 2020a) and suggest that different coronaviruses might have differential requirements or preferences for intracellular transport proteins.

We also uncovered an unexpected robust dependency on the Mediator complex (**Figure 3A, Figure S4B**). Intriguingly, a non-overlapping set of Mediator subunits was recently implicated in HIV-1 replication (Ruiz et al., 2014), suggesting that different viruses might have specific requirements for members of this complex during transcription and replication of their genomes. Therefore, one might speculate that selective targeting of host Mediator subunits using small molecules (e.g., CDK8 inhibitors; (Hofmann et al., 2020; Solum et al., 2020)) could be leveraged as a therapy to disrupt SARS-CoV-2 infection (Bancerek et al., 2013; Napoli et al., 2012; Spaeth et al., 2011).

Beyond well-characterized pathways represented by multiple components, we also identified factors with less understood network-level connections, including the DHX36 and DHX38 DEAH-Box helicases, the GOLGA6L1 and GOLGA8O Golgin family proteins, the TRMT5 and TRMT6 TRNA methyltransferases, the GPR89A and GPR89B G protein-coupled receptors, the TMED2 and TMED10 Transmembrane P24 trafficking proteins, and a pair of genes involved in phosphatidylethanolamine biosynthesis, such as *PCYT2* and *EPT1*, among others (**Figure 3A** and **Table S1**). Further mining of this SARS-CoV-2 host factor essentiality compendium could expand the repertoire of potential targetable factors to treat COVID-19.

The compendium of human factors co-opted by the HCoV-NL63 and HCoV-229E alphacoronaviruses significantly differs from that of the SARS-CoV-2 and HCoV-OC43 betacoronaviruses (**Figure 3A**). For instance, HCoV-NL63 seems particularly dependent on chromatin regulators with known functional interactions, including the KDM6A/UTX histone demethylase, the KMT2D/MLL4 histone methyltransferase, and the EP300 histone acetyltransferase, raising the interesting possibility that HCoV-NL63 co-opts the well established UTX-MLL4-EP300 enhancer remodeling network (Wang et al., 2017) to reprogram the transcriptome of the host for successful infection (**Figure 3A**). On the other hand, HCoV-229E seems particularly dependent on HOPS complex proteins, including VPS11 and RAB7A (**Figure 3A**) (Balderhaar and Ungermann, 2013). These results demonstrate that coronaviruses from the same genera, including the HCoV-NL63 and HCoV-229E alphacoronaviruses, can employ different host factor pathways during infection.

By integrating our *Coronaviridae* screening data with additional Zika virus and yellow fever virus CRISPR screening data (Hoffmann & Schneider et al.,; see accompanying manuscript), we identified TMEM41B as a critical host factor for infection by SARS-CoV-2, HCoV-OC43, HCoV-NL63, and HCoV-229E. TMEM41B is a poorly understood ER-localized transmembrane protein that was recently shown by three independent groups to regulate autophagy in conjunction with VMP1 (Moretti et al., 2018; Morita et al., 2018; Shoemaker et al., 2019). Strikingly, TMEM41B was the only autophagy-related gene that scored as a significant hit across the SARS-CoV-2, HCoV-OC43, and HCoV-229E CRISPR screens (**Figure 4A**), and it was subsequently validated as a co-factor for HCoV-NL63 as well (**Figure 4C**). This suggests a putative autophagy-independent role for TMEM41B as a pan-coronavirus replication factor. Moreover, our functional and mechanistic work showing that TMEM41B is a required host factor for infection by more than ten flaviviruses (Hoffmann & Schneider et al.,; see accompanying manuscript) suggested that TMEM41B is the only human protein that is critical for infection by both virus families. Thus, TMEM41B is a critical host factor required for infection by all of the coronaviruses tested in our study, as well as several other viruses of high public health interest, representing an attractive target for further investigation and development of antiviral therapies.

Coronavirus host factor discovery and validation is an active area of research with multiple studies appearing on preprint servers in recent months (Baggen et al., 2020; Heaton et al., 2020; Hoffmann et al., 2020a; Wang et al., 2020; Wei et al., 2020; Zhu et al., 2020b). Wei et al., performed genome-wide CRISPR screens in the African green monkey cell line Vero-E6 and reported strong dependency on ACE2 and CTSL, both of which were also identified in our study (Wei et al., 2020). Interestingly, while Wei et al., found that histone cell cycle regulator (HIRA) was a potent negative regulator of SARS-CoV-2 in Vero-E6 cells, our study found the opposite in Huh-7.5 cells. Heaton et al., performed a similar screen in A549 cells engineered to express ACE2 and identified serine/arginine-rich protein-specific kinase 1 (SRPK1) as a single dominant hit that is unique to their screen (Heaton et al., 2020). They showed that an SRPK1/2 inhibitor was inhibitory to HCoV-229E and SARS-CoV-2 at 17–53 µM *in vitro*, a concentration well above the Ki for SRPK1 with unknown consequences to cell viability. Like Heaton et al., Zhu et al., performed a similar screen using A549-ACE2 cells (Zhu et al., 2020b). However, in contrast to Heaton et al., they identified known host factors such as ACE2 and CTSL, as well as elements of the retromer complex, the COMMD/CCDC22/CCDC93 (CCC) complex, the Wiskott– Aldrich syndrome protein and SCAR homologue (WASH) complex, the actin-related proteins-2/3 (Arp2/3) complex and others, but not SRPK1. More recently, Wang et al., performed a series of screens using Huh7.5.1-ACE2-IRES-TMPRSS2 cells (Wang et al., 2020). Of note, Huh7.5.1 cells do not express ACE2 in contrast to the parental Huh-7.5 cells used in the present study. Wang et al., identified four significant host factors, one of which was SCAP, validating our prior finding that SCAP plays an important role in coronavirus infection (Hoffmann et al., 2020a). The remaining three significant host factors were also identified in our study: TMEM106B, VAC14, and ACE2. By contrast, our study identified 128 high-confidence host factors, perhaps in part due to our inclusion of multiple replicates per screen and two temperatures for SARS-CoV-2 as compared to single replicates and temperatures in the Wang et al., SARS-CoV-2 screen. Most recently, Baggen et al., reported results of SARS-CoV-2 and HCoV-229E screens using Huh-7 cells (Baggen et al., 2020). Their study highlighted TMEM106B, similar to Wang et al., and the present study, as well as TMEM41B, the focus of the present study and our accompanying paper on the role of TMEM41B as a pan-flavivirus host factor (Hoffmann & Schneider et al.,; see accompanying manuscript). Baggen at al., further demonstrated activity of phosphatidylinositol 3-kinase catalytic subunit type 3 (PIK3C3) / vacuolar protein sorting 34 (VPS34) inhibitors across multiple coronaviruses, although the mechanism of action for SARS-CoV-2 is unclear as PIK3C3 appeared to be dispensable for SARS-CoV-2 based on their results and on the present study, although it does appear essential for HCoV-229E and HCoV-OC43 in multiple studies described above.

The results of this study should be interpreted within the context of its limitations. If a gene did not score in our screens, it does not rule out that gene being an important SARS-CoV-2 factor for a number of reasons. First, pooled CRISPR screens may not identify functionally redundant or buffering genes (Ewen-Campen et al., 2017). Second, Huh-7.5 cells were chosen based on their infectivity by multiple coronaviruses; however, it should be understood that they are not airway cells. Nevertheless, a recent study demonstrated that hits in Huh-7 cells translate to human cells of lung origin (Baggen et al., 2020). Furthermore, as shown in **Figure S6**, the vast majority of genes identified here are expressed in human cells and tissues known to be infected by SARS-CoV-2. Lastly, our current experimental system is limited to assessing survival and can not interrogate host factors that act in late stages of the viral life cycle, nor can it identify genes that play important roles in immune modulation and pathogenesis.

It is essential to understand the underlying biology of a disease in order to develop new strategies for treatment and prevention. For infectious diseases, this entails studying the biology of the pathogen and the host. We identified complex, interconnected networks of coronavirus host factors and pathways that are essential for virus infection, nominating hundreds of new host proteins that represent liabilities for SARS-CoV-2 and potential opportunities for therapeutic intervention. This represents the most extensive functional catalog of host factors required for infection by SARS-CoV-2 and three seasonal coronaviruses (HCoV-229E, HCoV-NL63, and HCoV-OC43) to date, providing a larger context in which to interpret ongoing and future large scale CRISPR studies. Future efforts will focus on dissecting the complex interplay between virus and host and direct medicinal chemistry and drug repurposing resources toward the most chemically-tractable targets.

## ACKNOWLEDGEMENTS

This work was initiated and conducted under unusual circumstances. As New York City and much of the world was sheltering in place to reduce the spread of SARS-CoV-2, all of the authors here were sustained during the shutdown by generous funding intended for related and unrelated work. During this time we were fortunate to obtain funding from government and charitable agencies that allowed this COVID-19 work to continue. For funding directly related to these COVID-19 efforts, we thank the G. Harold and Leila Y. Mathers Charitable Foundation and the Bawd Foundation for their generous awards. We also thank Fast Grants (www.fastgrants.org), a part of Emergent Ventures at the Mercatus Center, George Mason University. Research reported in this publication was supported in part by the National Institute of Allergy and Infectious Diseases of the National Institutes of Health under Award Number R01AI091707. The content is solely the responsibility of the authors and does not necessarily represent the official views of the National Institutes of Health. We further received funding for COVID-19 related work from an administrative supplement to U19AI111825. As mentioned above, authors were also supported with non-COVID-19 funding by the following awards, foundations, and charitable trusts: R01CA190261, R01CA21344, U01CA213359, R01AI143295, R01AI150275, R01AI143295, R01AI116943, P01AI138938, P30CA008748, P30CA016087, R03AI141855, R21AI142010, W81XWH1910409, EMBO Fellowship ALTF 380-2018, F32AI133910, the Robertson Foundation, and an Agilent Technologies Thought Leader Award. S.W.L. is the Geoffrey Beene Chair of Cancer Biology and a Howard Hughes Medical Institute Investigator. F.J.S-R is a HHMI Hanna Gray Fellow and was partially supported by an MSKCC Translational Research Oncology Training Fellowship (NIH T32-CA160001). We also thank the NYU Langone Health Genome Technology Center. We also wish to thank Rodrigo Romero (MSKCC) for assistance with Revolve fluorescence microscopy and Aileen O’Connell, Santa Maria Pecoraro Di Vittorio, Glen Santiago, Mary Ellen Castillo, Arnella Webson, and Sonia Shirley for outstanding administrative or technical support.

## AUTHOR CONTRIBUTIONS

Conceptualization: WMS, HHH, JML, FJSR, AL, JTP

Methodology: WMS, HHH, JML, FJSR, AL, JTP

Formal analysis: JML, JTP

Investigation: WMS, HHH, AL, JLP, EM, IRL, AFS, JTP

Resources: SWL, CMR, JTP

Data curation: JML, JLP, JTP

Supervision: CMR, JTP

Visualization: JML, FSR, JTP

Writing - original draft: FJSR, JML, JTP

Writing – review & editing: WMS, JML, HHH, FJSR, JLP, IRL, MRM, CMR, JTP

Project Administration: AWA, AP, MRM

Funding acquisition: SWL, CMR, JTP

## LEAD CONTACT AND MATERIALS AVAILABILITY

Further information and requests for resources and reagents should be directed to and will be fulfilled by the Lead Contacts, John T. Poirier and Charles M. Rice.

## MATERIALS AND METHODS

### Plasmids and sgRNA cloning

To generate stable Cas9-expressing cell lines, we used lentiCas9-Blast (Addgene, cat. #52962). To express sgRNAs, we used lentiGuidePurov2, a variant of lentiGuide-Puro (Addgene, cat. #52963) that contains an improved sgRNA scaffold based on Chen et al. 2013 (Chen et al., 2013). For sgRNA cloning, lentiGuidePurov2 was linearized with BsmBI (NEB) and ligated with BsmBI-compatible annealed and phosphorylated oligos encoding sgRNAs using high concentration T4 DNA ligase (NEB). HIV-1 Gag-Pol and VSV-G plasmid sequences are available upon request.

### Cell culture

Lenti-X 293T™ cells (*H. sapiens*; sex: female) obtained from Takara (cat. #632180) and Huh-7.5 cells (*H. sapiens*; sex: male) (Blight et al., 2002) were maintained at 37 °C and 5% CO2 in Dulbecco’s Modified Eagle Medium (DMEM, Fisher Scientific, cat. #11995065) supplemented with 0.1 mM nonessential amino acids (NEAA, Fisher Scientific, cat. #11140076) and 10% hyclone fetal bovine serum (FBS, HyClone Laboratories, Lot. #AUJ35777). Both cell lines have tested negative for contamination with mycoplasma.

### Production and titration of coronavirus stocks

SARS-CoV-2 (strain: USA-WA1/2020) and HCoV-NL63 were obtained from BEI Resources (NR-52281 and NR-470). HCoV-OC43 was obtained from ZeptoMetrix (cat. #0810024CF) and HCoV-229E was generously provided by Volker Thiel (University of Bern). All viruses were amplified at 33 °C in Huh-7.5 cells to generate a P1 stock. To generate working stocks, Huh-7.5 cells were infected at a multiplicity of infection (MOI) of 0.01 plaque forming unit (PFU)/cell (SARS-CoV-2, HCoV-NL63, HCoV-OC43) and 0.1 PFU/cell (HCoV-229E) and incubated at 33 °C until virus-induced CPE was observed. Supernatants were subsequently harvested, clarified by centrifugation (3,000 *g* × 10 min) at 4 dpi (HCoV-229E), 6 dpi (SARS-CoV-2, HCoV-OC43) and 10 dpi (HCoV-NL63), and aliquots stored at −80 °C.

Viral titers were measured on Huh-7.5 cells by standard plaque assay. Briefly, 500 µL of serial 10-fold virus dilutions in Opti-MEM were used to infect 4 × 10^5^ cells seeded the day prior into wells of a 6-well plate. After 90 min adsorption, the virus inoculum was removed, and cells were overlaid with DMEM containing 10% FBS with 1.2% microcrystalline cellulose (Avicel). Cells were incubated for 4 days (HCoV-229E), 5 days (SARS-CoV-2, HCoV-OC43) and 6 days (HCoV-NL63) at 33 °C, followed by fixation with 7% formaldehyde and crystal violet staining for plaque enumeration. All SARS-CoV-2 experiments were performed in a biosafety level 3 laboratory.

To confirm the identity of the viruses, RNA from 200 µl of each viral stock was purified by adding 800 µl TRIzol™ Reagent (ThermoFisher Scientific, cat. #15596026) plus 200 µl chloroform then centrifuged at 12,000 *g* x 5 min. The upper aqueous phase was moved to a new tube and an equal volume of isopropanol was added. This was then added to an RNeasy mini kit column (Qiagen, cat. #74014) and further purified following the manufacturer’s instructions. Viral stocks were confirmed via next generation sequencing at the NYU Genome Technology Center using an Illumina stranded TruSeq kit and omitting the polyA selection step. Libraries were then sequenced by MiSeq Micro (2 × 250 bp paired end reads).

### Infection of TMEM41B knockout cells with SARS-CoV-2, HCoV-OC43, HCoV-NL63, and HCoV-229E

Huh-7.5 TMEM41B knockout cells (KO) and their reconstituted (tagRFP-TMEM41B) counterpart were generated as described in (Hoffmann & Schneider et al.,; see accompanying manuscript). The day prior to infection, Huh-7.5 WT, TMEM41B KO and reconstituted KO cells were seeded into 96-well plates at different densities relative to time of fixation e.g., 1×10^4^, 7.5×10^3^ and 5×10^3^ cells/well for a 24, 48, and 72 hours post infection time point, respectively. Cells were infected with the different coronaviruses under optimal conditions for each virus by directly applying 50 uL of virus inoculum to each well (n=3) at the following MOIs: SARS-CoV-2: MOI = 0.05 PFU/cell, HCoV-OC43: MOI = 2 PFU/cell, HCoV-NL63: MOI = 0.05 PFU/cell, HCoV-229E: MOI = 0.15 PFU/cell, and incubated for 24 hours at 37 °C (HCoV-229E), 48 hours at 33 °C (SARS-CoV-2), and 72 hours at 33 °C (HCoV-OC43 and HCoV-NL63). Cells were subsequently fixed by adding an equal volume of 7% formaldehyde to the wells followed by permeabilization with 0.1% Triton X-100 for 10 minutes. After extensive washing, cells were incubated for 1 hour at room temperature with blocking solution of 5% goat serum in PBS (Jackson ImmunoResearch: catalog no. 005–000-121). To stain for SARS-CoV-2, a rabbit polyclonal anti-SARS-CoV-2 nucleocapsid antibody (GeneTex: catalog no. GTX135357) was added to the cells at a 1:1,000 dilution in blocking solution and incubated at 4 °C overnight. To detect infected cells for HCoV-229E, HCoV-OC43 and HCoV-NL63, a mouse monoclonal anti-dsRNA antibody (Scicons: catalog no. 10010500) was used under similar conditions. Goat anti-rabbit AlexaFluor 488 (Life Technologies: catalog no. A-11012) and goat anti-mouse AlexaFluor 488 (Life Technologies: catalog no. A-11001) were used as a secondary antibody at a dilution of 1:2,000. Nuclei were stained with Hoechst 33342 (ThermoFisher Scientific: catalog no. 62249) at a 1:1,000 dilution. Images for quantification of virus infection and cell viability were acquired with a fluorescence microscope and analyzed using ImageXpress Micro XLS (Molecular Devices, Sunnyvale, CA). Images for assessment of tagRFP-TMEM41B subcellular localization were obtained using a Revolve inverted microscope (Echo, San Diego, CA).

### CRISPR-Cas9 genetic screening

Huh-7.5-Cas9 cells were generated by lentiviral transduction of lentiCas9-Blast (Addgene, cat. #52962) followed by selection and expansion in the presence of 5 µg/ml blasticidin. The human CRISPR Brunello library (Doench et al., 2016b) was obtained through Addgene as a ready-to-use lentiviral pooled library at a titer ≥ 1×10^7^ TU/mL (Addgene, cat. #73178-LV). To deliver the Brunello sgRNA library, 2.04 × 10^8^ Huh-7.5-Cas9 cells were transduced by spinoculation at 1,000 *g* x 1 h in media containing 4 µg/ml polybrene (Millipore, cat. #TR-1003-G) and 20 mM HEPES (Gibco, cat. #15630080) at a MOI = 0.21 to achieve ∼560-fold overrepresentation of each sgRNA. Cells were spinoculated at 3 × 10^6^ cells/well in 12-well plates in 1.5 ml final volume. Six hours post transduction, cells were trypsinized and transferred to T175 flasks at 7 × 10^6^ cells/flask. Two days later, media was replaced with fresh media containing 1.5 µg/ml puromycin and cells were expanded for five additional days prior to seeding for coronavirus infection. Huh-7.5-Cas9 cells transduced with the Brunello sgRNA library were seeded in p150 plates at 4.5 × 10^6^ cells/plate with two plates per replicate (e.g., 9 × 10^6^ cells) and three replicates for each condition (mock, HCoV-229E, HCoV-NL63, HCoV-OC43).

For biosafety reasons, SARS-CoV-2 infection was performed in T175 screw top flasks. For infections at 37 °C, we seeded cells at 5 × 10^6^ cells per flask and used two flasks (e.g., 1 × 10^7^ cells) per replicate. For infections at 33 °C, we seeded cells at 6.7 × 10^6^ cells per flask and used three flasks (e.g., 2 × 10^7^ cells) per replicate. Both SARS-CoV-2 screens were performed in triplicate. The following day, the media was removed and viruses diluted in 10 ml/plate OptiMEM were added to cells. The inocula of HCoV-229E, HCoV-NL63 and SARS-CoV-2 were supplemented with 1 µg/ml TPCK-treated trypsin (Sigma-Aldrich, cat. #T1426) increasing the rate of infection. After two hours on a plate rocker at 37 °C, 10 ml/plate media was added and plates were moved to 5% CO2 incubators set to 33 °C (HCoV-229E, HCoV-NL63, HCoV-OC43, and SARS-CoV-2) or 37 °C (SARS-CoV-2). Coronavirus screens were performed at the following MOIs in PFU/cell: HCoV-229E = 0.05 at 33 °C; HCoV-NL63 = 0.01 at 33 °C; HCoV-OC43 = 1 at 33 °C; SARS-CoV-2 = 0.01 at 33 °C and 0.1 at 37 °C. Mock cells cultured at both temperatures were passaged every 3-4 days and re-seeded at 4.5 × 10^6^ cells/plate with two plates per replicate. Media was changed on virus infected plates as needed to remove cellular debris. Mock cells and cells that survived coronavirus infection were harvested approximately one to two weeks post infection.

Genomic DNA (gDNA) was isolated via ammonium acetate salt precipitation if greater than 1.5 × 10^6^ cells were recovered or using the Monarch Genomic DNA Purification kit (NEB) if fewer per the manufacturer’s instructions. gDNA concentrations were quantitated via UV spectroscopy and normalized to 250 ng/µl with 10 mM Tris. The library was amplified from gDNA by a two-stage PCR approach. For PCR1 amplification, gDNA samples were divided into 50 ul PCR reactions. Each well consisted of 25 µl of NEB Q5 High-Fidelity 2X Master Mix, 2.5 ul of 10 µM forward primer Nuc-PCR1_Nextera-Fwd Mix, 2.5 ul of 10 µM reverse primer Nuc-PCR1_Nextera-Rev Mix and 20 µl of gDNA (5 µg each reaction). PCR1 cycling settings: initial 30 s denaturation at 98 °C; then 10 s at 98 °C, 30 s at 65 °C, 30 s at 72 °C for 25 cycles; followed by 2 min extension at 72 °C. PCR1 samples were cleaned up by isopropanol precipitation, and normalized to 20 ng/µl. Each PCR2 reaction consisted of 25 µl of NEB Q5 High-Fidelity 2X Master Mix, 2.5 µl 10 µM Common_PCR2_Fwd primer, and 2.5 ul of 10 µM reverse i7 indexing primer. PCR2 cycling settings: initial 30 s at 98 °C; then 10 s at 98 °C, 30 s at 65 °C, 30 s at 72 °C for 13 cycles. PCR products were again purified by SPRI, pooled and sequenced on an Illumina NextSeq 500 at the NYU Genome Technology Center using standard Nextera sequencing primers and 75 cycles.

### Analysis of CRISPR-Cas9 genetic screen data

FASTQ files were processed and trimmed to retrieve sgRNA target sequences followed by enumeration of sgRNAs in the reference sgRNA library file using MAGeCK (Li et al., 2014). MAGeCK was also used to determine gene essentiality (beta) using its maximum likelihood estimation (MLE) algorithm. Z-scores for visualization in the form of heatmaps were computed using the following approach: for each condition, the log2 fold change with respect to the initial condition was computed. A natural cubic spline with 4 degrees of freedom was fit to each pair of infected and control cells and residuals were extracted. To obtain gene-wise data, the mean residuals for each group of sgRNAs was calculated, a z-score was computed, and a p-value was determined using a 2-sided normal distribution test. P-values were combined across screens using Fisher’s sumlog approach and corrected for multiple testing using the method of Benjamini & Hochberg.

### Functional clustering and network analysis of screening data

High confidence CRISPR hits with FDR cutoffs below 0.05 were extracted for functional clustering and network building. Briefly, enriched pathways were identified from the NIH NCATS BioPlanet database (Huang et al., 2019), which aggregates currates pathways from multiple sources, using competitive gene set testing of z scores in pre-ranked mode (Wu and Smyth, 2012). For construction of the network in Figure 3, significant CRISPR hits from any virus were searched using the STRING database (string-db.org) (Szklarczyk et al., 2019) using default parameters and imported into Cytoscape (Shannon et al., 2003). Overlapping hits per virus were calculated and subsequently depicted as pie charts per node in Adobe Illustrator. For virus specific networks in Figure S2, significant CRISPR hits per virus and the next adjacent 100 interactors were extracted and graphed in Cytoscape.

### Analysis of scRNAseq data

For scRNAseq analysis, Seurat objects were downloaded from FigShare (https://doi.org/10.6084/m9.figshare.12436517) (Chua et al., 2020). Dotplots for select cell identities and for all hgh confidence CRISPR hits per virus were plotted using the DotPlot function in Seurat (Stuart et al., 2019).

### Statistical Analyses

Statistical tests were used as indicated in the figure legends. Generation of plots and statistical analyses were performed using the R statistical computing environment. Error bars represent standard deviation, unless otherwise noted. We used Student’s *t*-test (unpaired, two-tailed) to assess significance between treatment and control groups, and to calculate P values. P < 0.05 was considered statistically significant.

## DATA AND CODE AVAILABILITY

Data supporting the findings of this study are reported in Supplementary Figures S1-S6 and Tables S1-S2. All raw data corresponding to CRISPR screens will be available through NCBI GEO. Networks are available on NDEx. All reagents and materials generated in this study will be available to the scientific community through Addgene and/or MTAs.

## SUPPLEMENTARY FIGURE TITLES AND LEGENDS

**Supplementary Figure S1.**
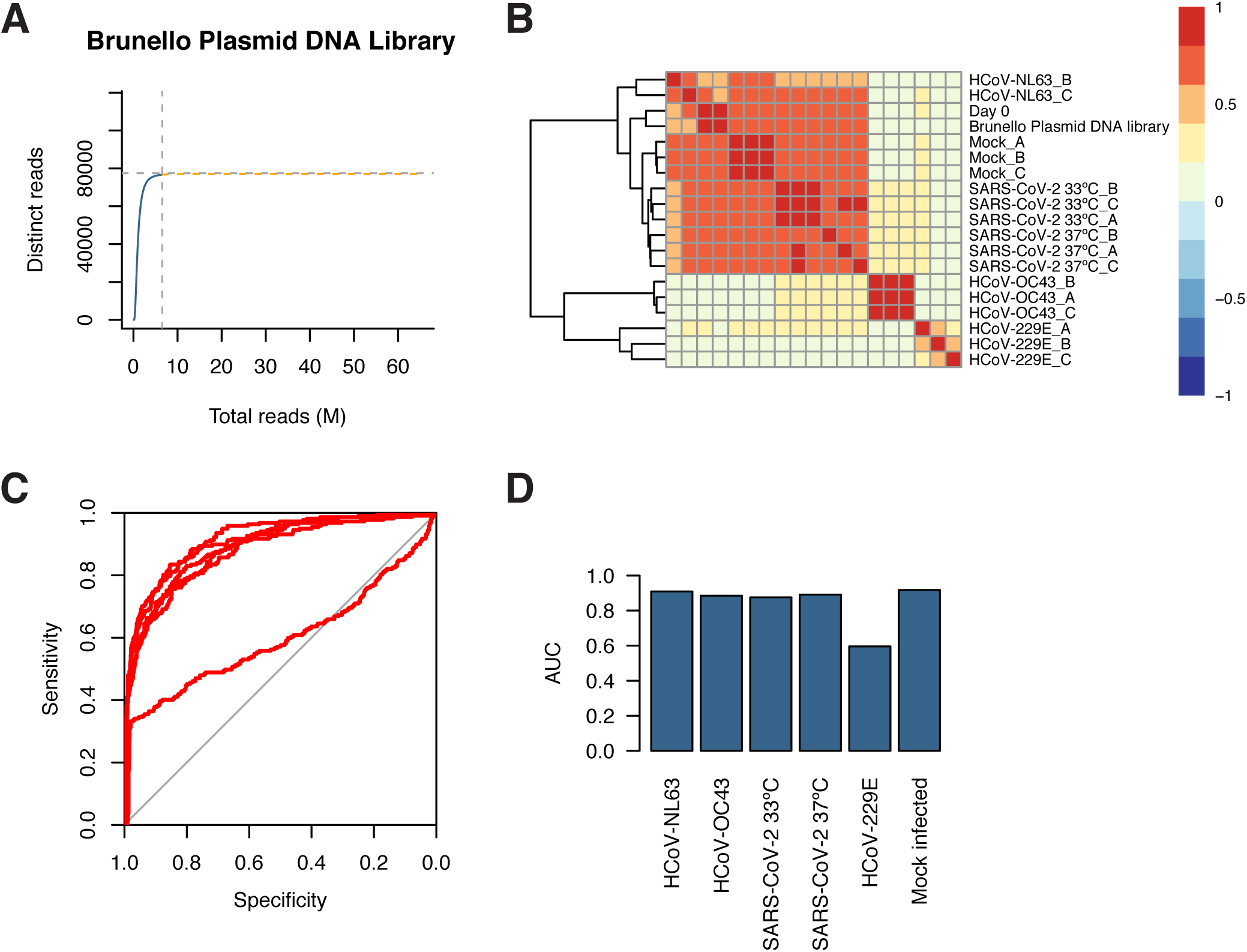
Screen quality control data. (A) Species accumulation curve for unique reads with ≥10 counts. (B) Heatmap for correlation coefficients between samples. (C) ROC curves for each screen. (D) Area under the curve (AUC) for each ROC curve in (C).

**Supplementary Figure S2.**
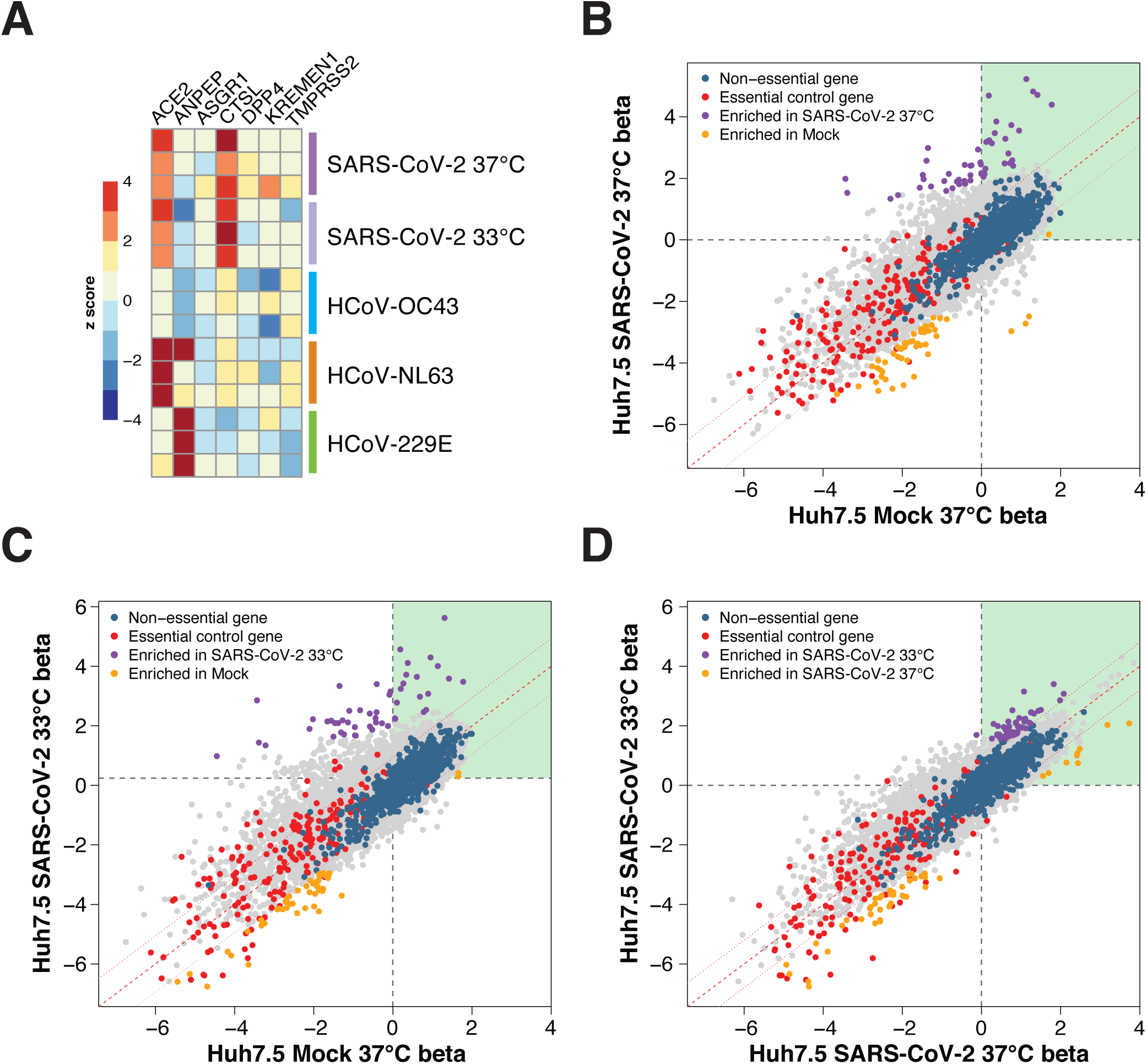
Analysis of established and putative coronavirus host factors and gene-wise fitness scores for SARS-CoV-2. (A) Heatmap of z scores for known and putative coronavirus host factors. (B-D) Genewise fitness beta scatterplots comparing SARS-CoV-2 at 37°C vs. mock (B), SARS-CoV-2 at 33°C vs. mock (C), and SARS-CoV-2 at 33°C vs. SARS-CoV-2 at 37°C (D). Non-targeting controls and essential control genes are highlighted in blue and red, respectively.

**Supplementary Figure S3.**
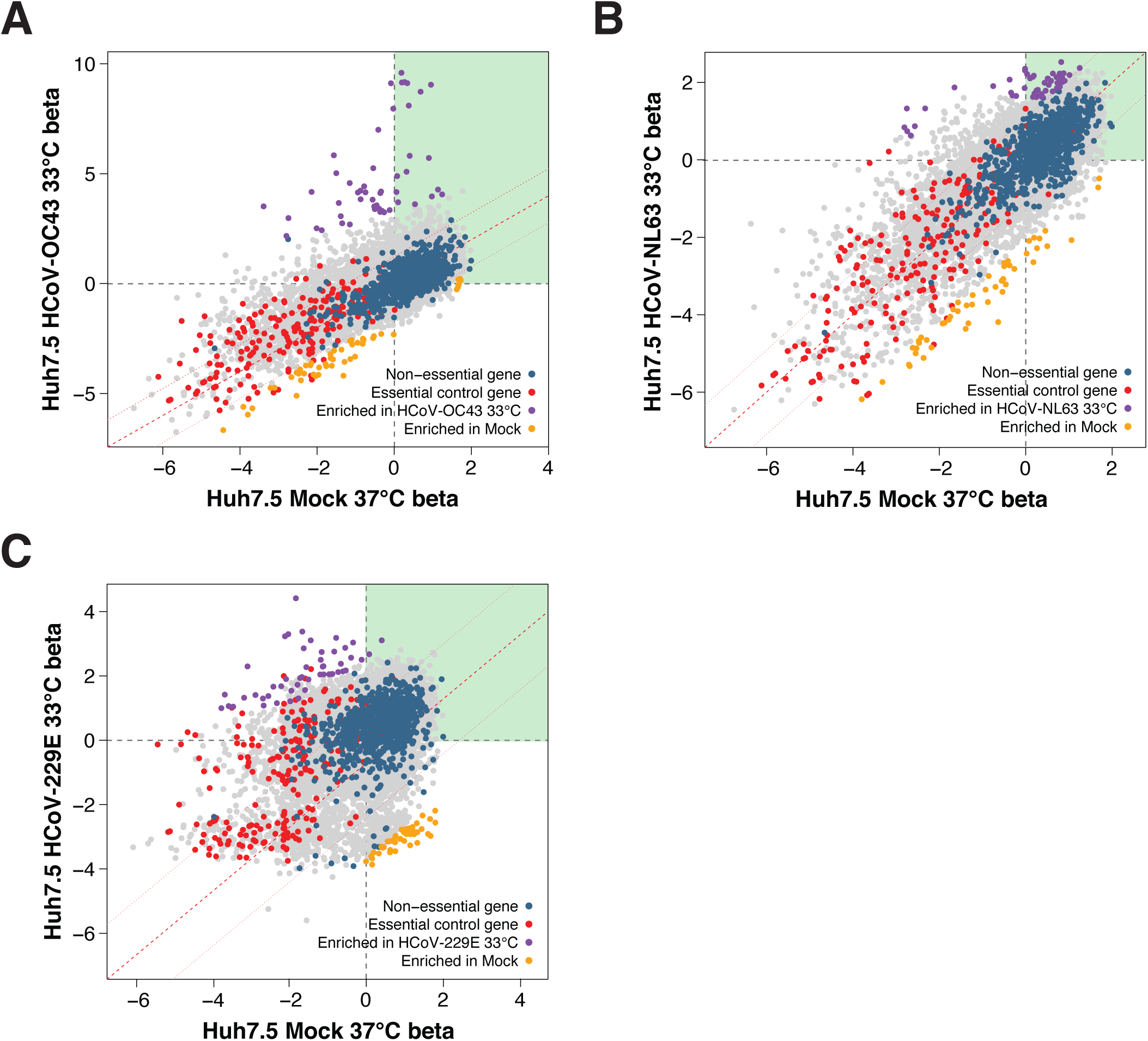
Gene-wise fitness scores for 3 seasonal coronaviruses. (A) HCoV-OC43 vs. mock infected. (B) HCoV-NL63 vs. mock infected. (C) HCoV-229E vs. mock infected.

**Supplementary Figure S4.**
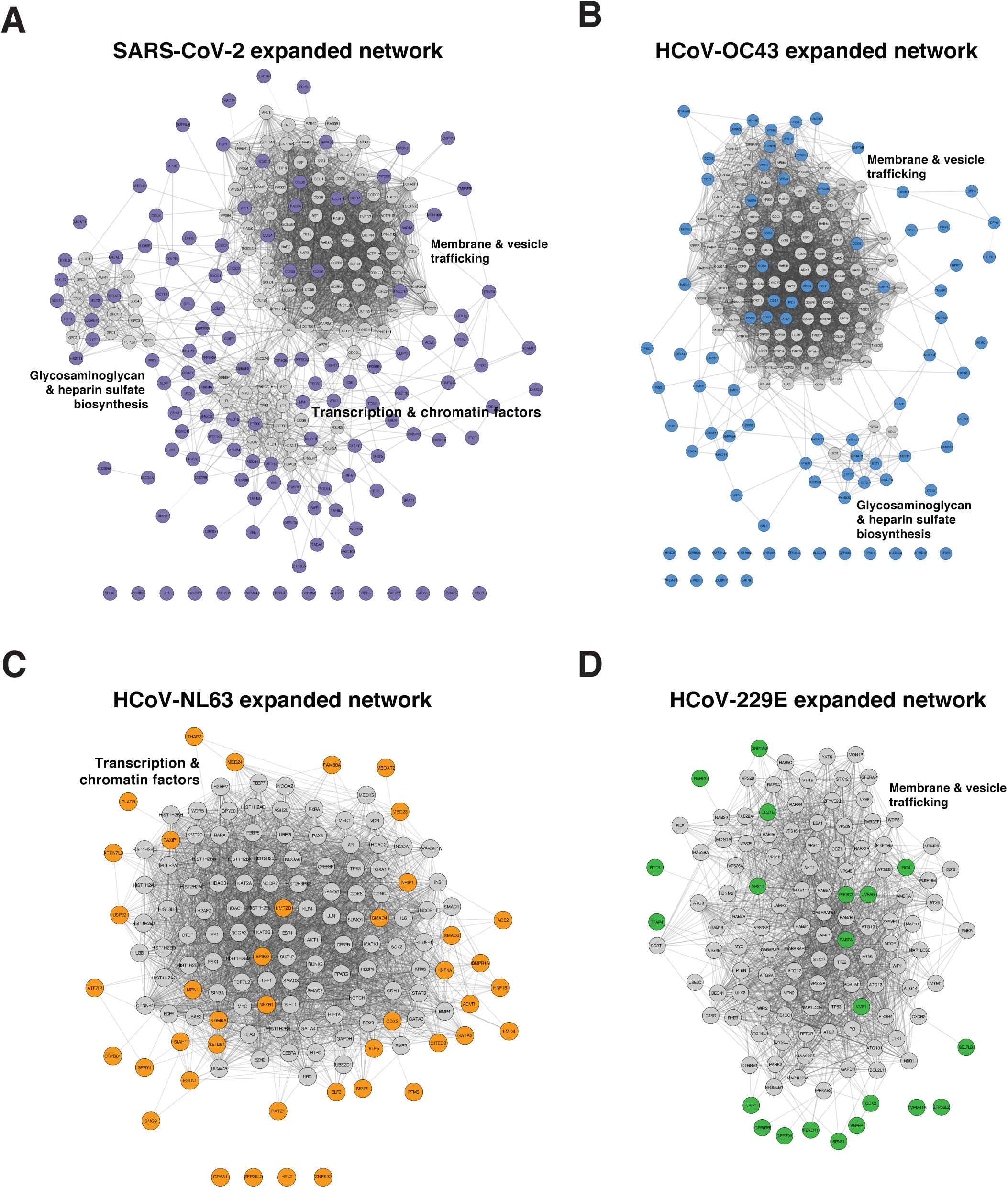
Expanded coronavirus specific networks. (A-D) Network diagram of all coronavirus screen hits and the next 100 adjacent interactors, shown in gray. Broad functional categories of highly interconnected gene neighborhoods are indicated. Diagrams are for SARS-CoV-2 (A), HCoV-OC43 (B), HCoV-NL63 (C) and HCoV-229E (D).

**Supplementary Figure S5.**
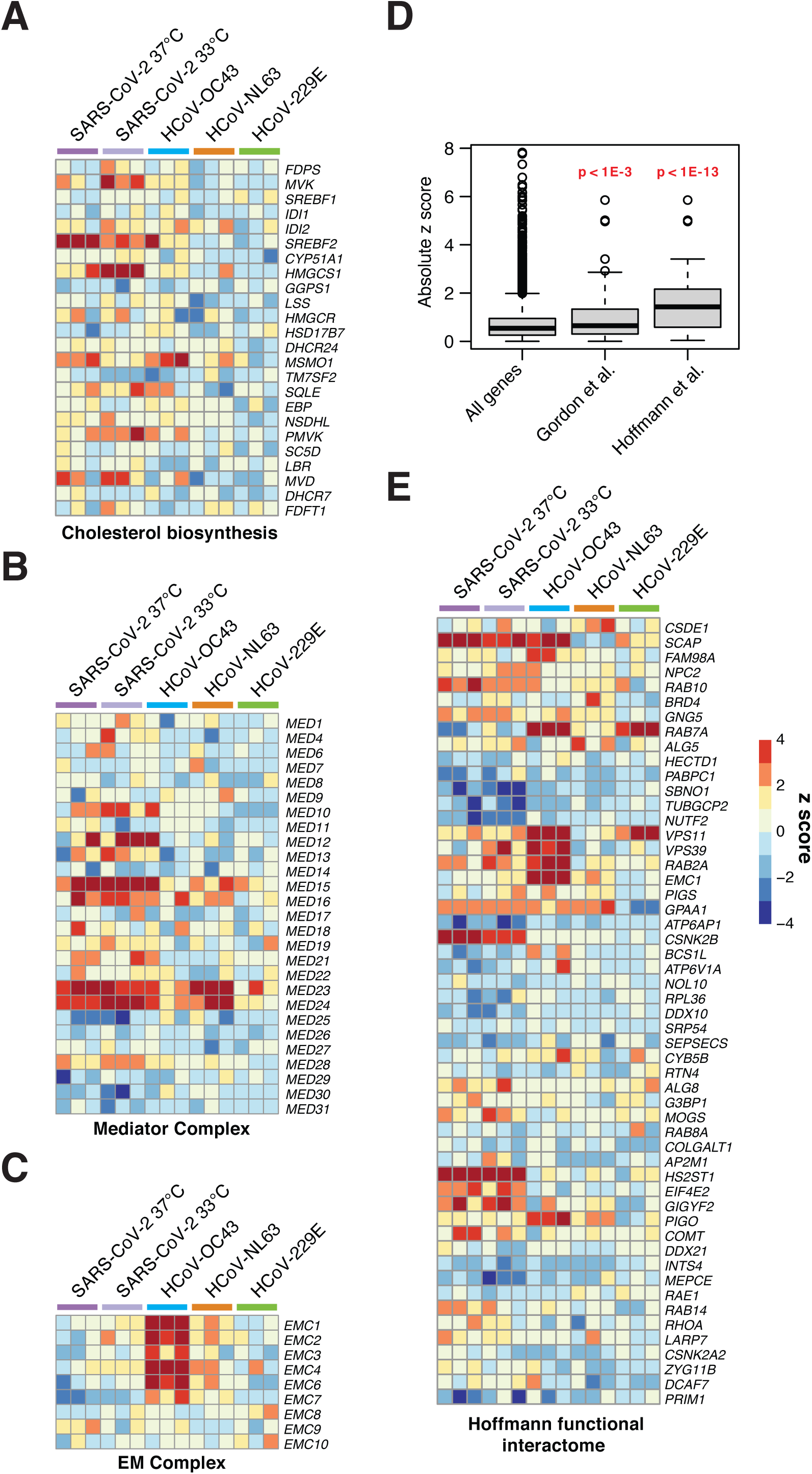
Additional enriched pathways and subset analysis of a high-confidence SARS-CoV-2 protein:protein interactome. (A) Heatmap of z scores for the cholesterol biosynthesis gene set. (B) Heatmap of z scores for the mediator complex gene set. (C) Heatmap of z scores for the endoplasmic reticulum membrane complex gene set. (D) Subset analysis of absolute z-scores for SARS-CoV-2 screen at 37 °C for the Gordon et al. high-confidence interactome and the Hoffmann et al., functional interactome. Significance tests are two-sided Wilcoxon rank sum test. (E) Heatmap of z scores from functional interactome hits in Hoffmann et al. subsetted from genome-scale CRISPR screens.

**Supplementary Figure S6.**
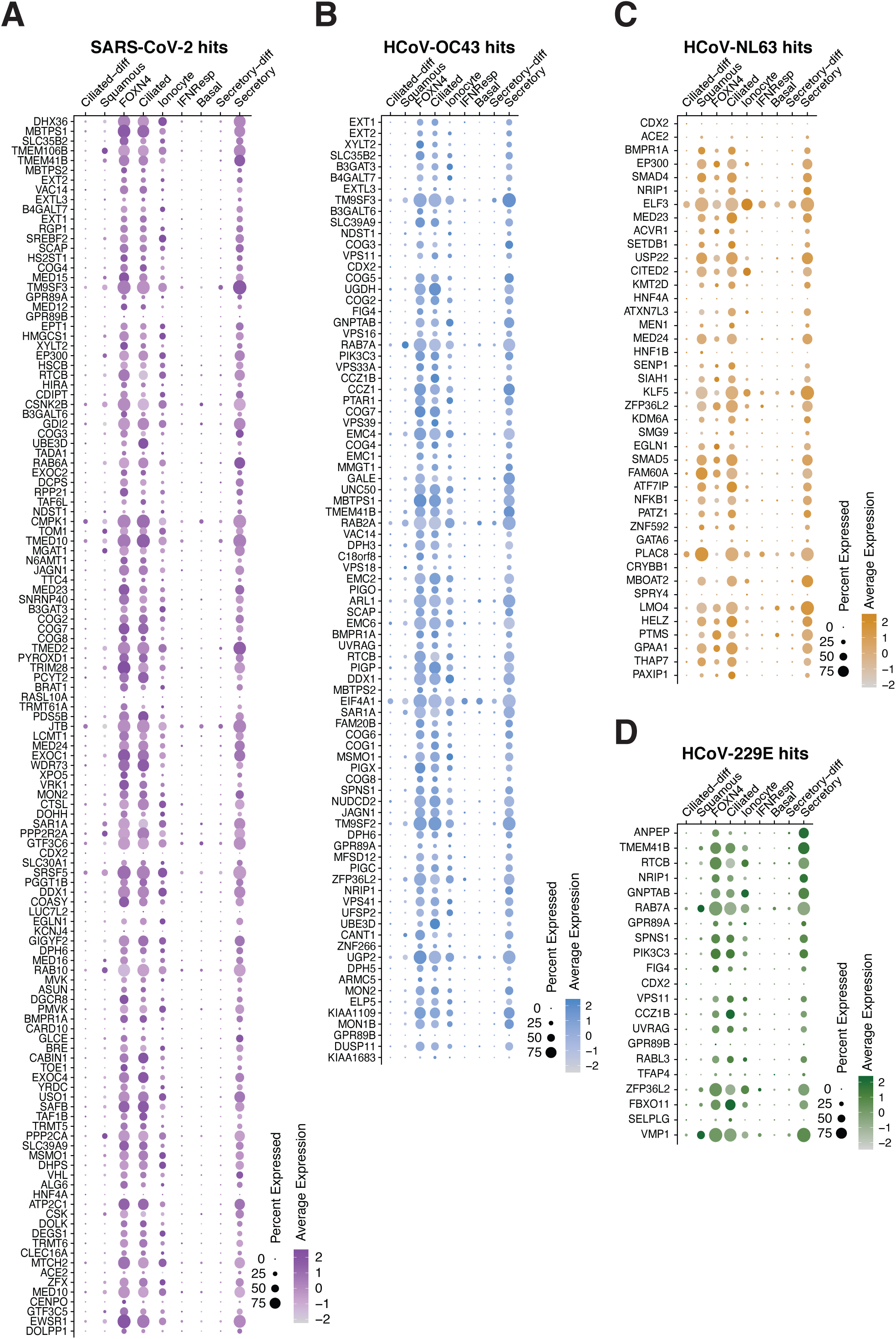
A majority of functional coronavirus host factors are expressed in the airway. (A-D) scRNAseq expression dotplot diagrams from select cells in the airway of the average expression and the percent of cells expressing coronavirus host factors for SARS-CoV-2 (A), HCoV-OC43 (B), HCoV-NL63 (C) and HCoV-229E (D). Rows for each diagram are ordered top to bottom from highest to lowest z-score. Data are from (Chua et al., 2020).

**Table S1. CRISPR screen results**.

Table S1A: Huh-7.5 37 °C SARS-CoV-2 z scores.

Table S1B: Huh-7.5 33 °C SARS-CoV-2 z scores.

Table S1C: Huh-7.5 33 °C HCoV-OC43 z scores.

Table S1D: Huh-7.5 33 °C HCoV-NL63 z scores.

Table S1E: Huh-7.5 33 °C HCoV-229E z scores.

**Table S2. Sequences of sgRNAs and primers**.

Table S2A: Primers used to prepare Illumina-compatible libraries for high-throughput sequencing of CRISPR screens.

Table S2B: Full sequence of sgRNA expression lentiviral vectors constructed and used in this study.

